# Event-marked Windowed Communication: Inferring activity propagation from neural time series

**DOI:** 10.1101/2024.07.30.605466

**Authors:** Varun Madan Mohan, Thomas F. Varley, Robin F. H. Cash, Caio Seguin, Andrew Zalesky

## Abstract

Tracking signal propagation in nervous systems is crucial to our understanding of brain function and information processing. Current methods for inferring neural communication track patterns of sustained co-activation over time, making them unsuitable to detect discrete instances of signal transmission. Here, we propose Event-marked Windowed Communication (EWC), a new analytical framework to infer functional interactions arising from discrete signalling events between neural elements, in otherwise continuous time series data. In contrast to conventional measures of functional connectivity, our method utilises an event-based subsampling of neural time series, which allows it to capture the statistical analogue of activity propagation. We test EWC on simulations of neural dynamics and show that it is capable of retrieving ground truth motifs of directional signalling, over a range of model configurations. Critically, we demonstrate that EWC’s subsampling approach affords profound reductions in computation times, compared to established network inference methods such as transfer entropy. Lastly, we showcase the utility of EWC to infer whole-brain functional networks from MEG recordings. Networks computed using EWC and transfer entropy were highly correlated (median r=0.821 across subjects), but EWC inference was approximately 6.5 times faster per epoch. In summary, our work presents a new method to infer signalling from time series of neural activity at low computational costs. Our framework is flexible and can be applied to activity time series captured by diverse functional neuroimaging modalities, opening up new avenues for the study of neural communication.

## Introduction

Communication between neural elements — neurons, neural populations and grey matter regions, plays a central role in the functioning of the brain. Understanding the principles by which signals are dynamically and flexibly transmitted in networked nervous systems remains an open challenge and active area of research^1–3^. Efforts in this direction have produced a vast number of models of brain communication^1,4,5^, ranging from information transfer via neural oscillations^6–10^ to network measures of connectome communication^11–13^. Despite the abundance in models, efforts in model validation have been lacking, and it remains unclear which methods faithfully describe empirical patterns of neural dynamics and signalling. A key barrier to progress in model validation is our current inability to infer events of signal transmission from recordings of neural activity.

Functional connectivity (FC) is commonly used as a proxy for neural communication. Broadly defined, FC quantifies statistical dependencies in time series of neural activity using measures such as the Pearson correlation or Mutual Information, capturing the extent to which the dynamics of two brain regions are synchronised over a period of time. FC estimation has been extensively used in model formulation and validation as well as in experimental and clinical studies^3,14–24^. While useful, these FC measures are limited to capturing stable, sustained statistical associations over time, which can dilute or mask discrete events of directional signal transmission from one neural element to another. Information-theoretical measures such as Transfer Entropy stand to address this issue, but are typically data- and computation-heavy for whole-brain network inference^25^.

Explicitly tracking how activity (endogenous/exogenous perturbations) or signals propagate across the underlying anatomical substrate can provide important insights into the principles of inter-areal communication and help validate existing computational models. Current limitations in neuroimaging technology have seen activity propagation tracing often applied to smaller controlled stimulation experiments in microscale neuronal networks ^26–28^. Its application at the whole-brain scale is still limited^29–32^, although approaches combining existing data to yield robust insights seem to show promise^33^. In this work, our aim was to develop an efficient method of inferring activity propagation patterns by combining the practicality and efficiency of undirected FC estimation with the dynamical resolution afforded by activity propagation tracing. We term this modified FC estimation protocol Event-marked Windowed Communication (EWC). Specifically, EWC involves identifying salient events in neural recordings (“stimuli”) and the subsequent estimation of FC within short temporally ordered windows/subsamples of the signal. In this manner, EWC captures an indirect statistical analogue of activity propagation as opposed to spatiotemporally localised stimulation-response measurements. We first validate and demonstrate the utility of the EWC implementation using a simple in-silico network motif with ground truth signalling embedded in its dynamics. We also show that the EWC approach allows us to use undirected FC measures to capture directional interactions. We then study the computational tractability of the approach by comparing network inference runtimes as a function of number of nodes. Finally, we demonstrate a real-world application of the method on source-localised resting-state magnetoencephalography (MEG) recordings.

## Results

In this study, we develop a new method to quantify functional interactions between neural elements based on the detection of directional signalling events in their activity time series (Fig.1). The Event-marked Windowed Communication (EWC) protocol is composed of 3 main elements: 1) Event-identification 2) Temporal ordering based on conduction delays, and 3) Windowing or subsampling. Specifically, our approach involves first identifying significant deviations from the mean of activity time series, which we term “supra-threshold events” (For brevity, we will refer to them as “events” for the remainder of this paper). We then trace the downstream effects of these events on the activities of other neural elements, while considering the possible delays in signal conduction due to inter-element distances. Importantly, we restrict the estimation of communication to a short “communication window”, that starts at the timepoint of an event, for the source, and at a future timepoint for the target (proportional to the source-target distance). Once a communication window is specified, the magnitude of the signalling event is quantified by the statistical association between the within-window activity time series of the source and target. EWC is flexible and this association can be computed using various measures of FC, such as the partial correlation or Mutual Information. Lastly, we derive an aggregate communication measure by averaging the FC across communication windows.

**Figure 1.**
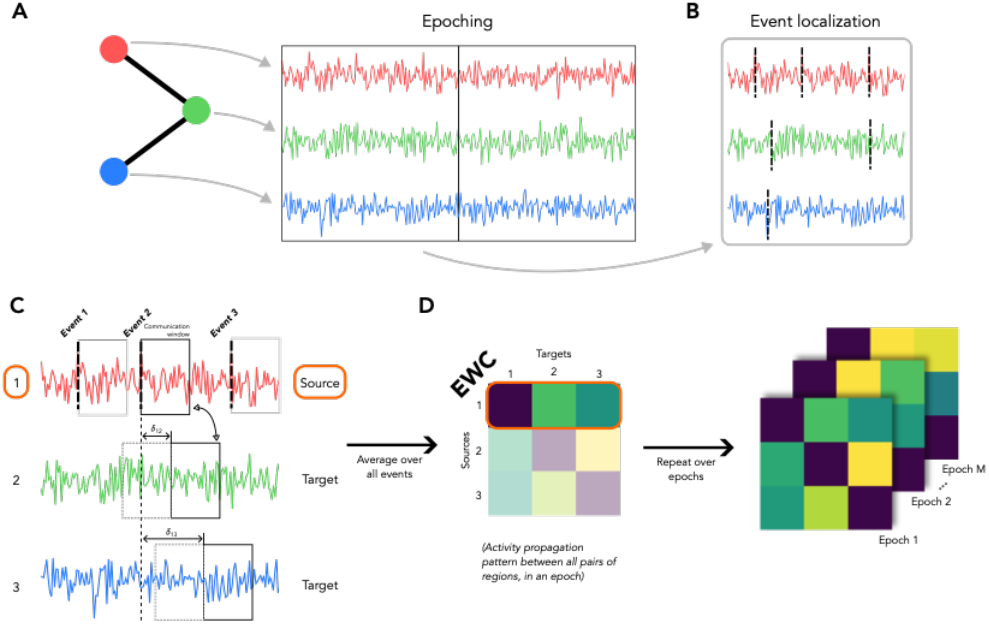
A pipeline to estimate communication patterns from regional time series. (A) Regional activities are divided into 10 second epochs, and each epoch is processed independently. (B) For each epoch, the regional activities are z-scored to identify instances of deviation from mean behaviour, which are termed “supra-threshold events”. (C) Each region is successively chosen as a “source”. For each event of a source, a communication window that encompasses the activity over a duration of 1 second, is defined starting at the time point of the event. Similar communication windows are defined over the activities of all other regions (termed “targets”), starting at timepoints delayed with respect to the source event, in proportion to the Euclidean distance between the source and target. (D) For each event for a chosen source, the Event-marked Windowed Communication (EWC) is estimated between the source and all possible targets by computing the conditional mutual information between the activities contained in the communication windows of the source and target. The activity of the target over 1 second in the past is used as the conditioning variable. The pairwise EWC values over all events of a source (within an epoch) are averaged, to populate a row of the EWC matrix corresponding to the source index. Steps B-D are repeated over all epochs to give epoch-level EWC matrices that capture the dynamic communication patterns over the entire scan duration.

The analyses of this paper can be divided into three main sections. First, we tested our method in simulated time series produced by a simple connectivity motif in which node activity was governed by Linear Stochastic Model (LSM) dynamics (Fig 2A). The simple dynamical landscape of the LSM allowed us to impose ground truth signalling between nodes. We assessed our method’s ability to identify these communication patterns in relation to previously proposed measures of FC. Second, we characterise gains in computational time afforded by estimating FC using EWC. Third, we apply our method on source localised MEG recordings, and gauge the agreement between the FC inferred via EWC and bivariate Transfer Entropy.

**Figure 2.**
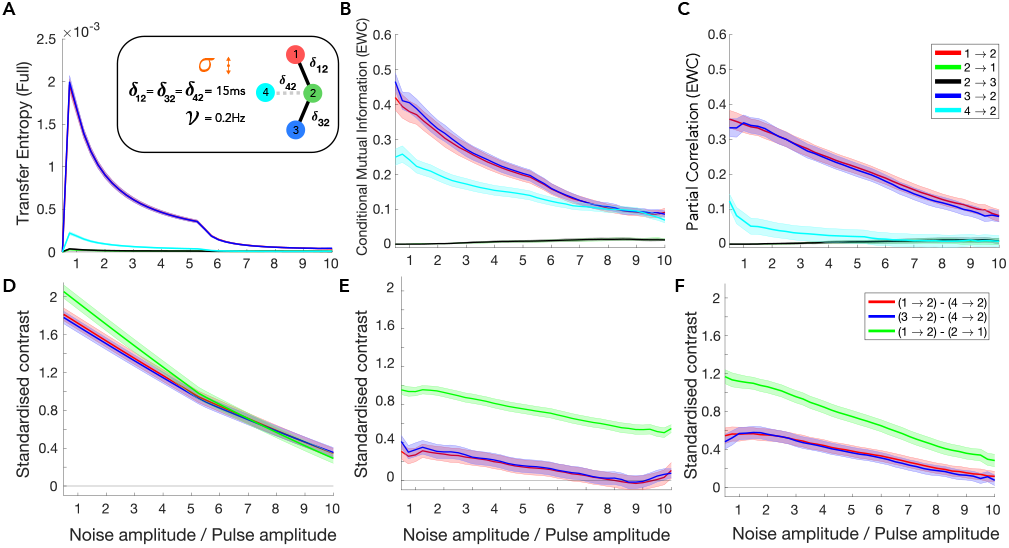
Communication over a network motif. (A) (inset) We test our method of estimating communication patterns on a simple 3-node network motif with a linear topology and an isolated node (4-cyan). The activities of the individual nodes in this network are described by a Linear Stochastic Model. The dynamics of the red, blue and cyan nodes have an additional Poisson process, causing it to spike at an average rate of 0.2 Hz, emulating communication events. δ_xy_ is the delay between nodes x and y, in ms. The noise amplitude of the LSM, α, is varied relative to the fixed Poisson pulse amplitude of the sources. (A) Transfer entropy (in nats) between nodes, estimated over the full signal. (B) Conditional Mutual Information (in nats) between nodes, estimated as per the EWC protocol (C) Partial Correlation (Pearson-R) between nodes, estimated as per EWC. We show only the absolute Pearson correlation. (D-F) Standardised contrast (difference between EWC of True and False connections, normalised by their pooled standard deviation) plots associated with the FC estimates. A non-zero contrast value indicates discriminability. Shading in the plots represent ±SEM (20 trials). All plots maximally smoothed (i.e. using all available points) to clearly reveal trends.

In the remainder of the paper, we will use the suffix “-Full” after the FC measure to indicate a conventional estimation technique using the entire signal, and “-EWC” to indicate our implementation.

### Asymmetric signalling over a network motif

We tested the EWC protocol in a four-node motif with Linear Stochastic Model (LSM) dynamics (Fig.3). Three of the nodes in this network were connected in a linear topology (nodes 1, 2, and 3), whereas node 4 was isolated (i.e. not influenced by the dynamics of other nodes). In addition to the LSM dynamics, the nodes 1, 3, and 4 also pulsated at random times according to a Poisson process. These pulses emulated punctual events of directional signalling, with known sources at nodes 1, 3, and 4.

**Figure 3.**
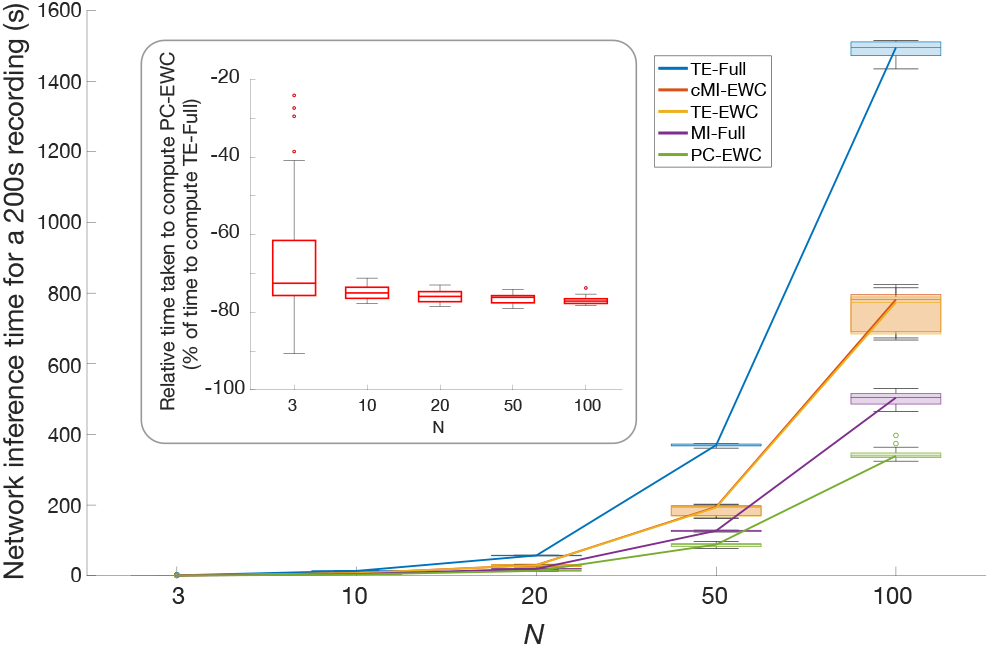
Time taken for network inference as a function of network size. Signals spanning 200s (Twenty 10s epochs) sampled from source-localised MEG recordings of 3 subjects. Sampling repeated 10 times per subject for each network size (Each box plot spanning 30 data points). Circles represent outliers. (Inset) Percentage of time taken to compute PC-EWC, relative to the time taken to compute TE-Full. For a range of network sizes, PC-EWC takes ≈75% less time to compute for a 200s scan, or equivalently, is ≈4 times faster.

EWC was implemented using two measures of within-window FC, partial correlation (PC-EWC) and conditional mutual information (cMI-EWC). Note that, traditionally, these are symmetric measures that cannot resolve directionality of functional interactions. We benchmarked EWC against Transfer Entropy (TE-Full), which was conventionally estimated based on the entire time series. We compared the ability of these measures to retrieve the ground truth signalling motif in simulations with increasing noise levels. To enable comparison of the different measures, we estimated the standardised contrast between the FC in true (existing) and false (absent) connections.

We considered the performance of our method as a function of the ratio between the noise level in the system and the amplitude at which the source regions pulsate. The delay between the sources (nodes 1, 3, and 4) and the target (node 2) was set at 15ms, and the sources pulsated according to a Poisson process with a mean frequency of 0.2Hz (Fig.2A - inset). The ground truth comprises connections from node 1 to node 2 and 3 to 2. We observed that estimating the TE-Full resulted in a good representation of the ground truth, with strong interactions from nodes 1 and 3 to 2 (Fig.2A). Importantly, the corresponding contrast distribution indicated that the interaction between node 4 (isolated) and 2 could be clearly discriminated from the true interactions, providing evidence of good specificity (Fig.2D). When the FC was instead measured using cMI-EWC, we observed an interaction between regions 4 and 2, although this false positive interaction was considerably weaker than the ground truth interactions from node 1 to 2, and 3 to 2, particularly at low noise levels (Fig.2B). Interestingly, PC-EWC performed well when compared to cMI-EWC. Like TE-Full, PC-EWC also resulted in a good representation of the ground truth (Fig.2C). The contrast between estimated interactions showed that true connections could be distinguished from absent connections even at large levels of noise (Fig.2F), although not as well as TE-Full. Importantly, in the case of cMI-EWC and PC-EWC, asymmetric interactions were estimated, despite the FC measures being undirected/symmetric. This asymmetry is a result of the temporal ordering within the EWC implementation.

Additionally, we also tested whether a common source equidistant from two targets would result in a spurious functional link between them (a closed triangle problem). For this we set region 2 as the pulsating source and estimated the PC-EWC. We observed that our method correctly identified communication from node 2 to both 1 and 3, with negligible communication in the opposite direction, or between the targets (Fig.S1). We also tested the EWC implementation over a range of conduction delays, and mean firing rates of the sources (Fig.S2).

To summarise, our comparative analyses demonstrate that EWC can accurately capture directional interactions over a range of noise levels, delays, and firing rates. We emphasize that PC and cMI, being undirected/symmetric measures, were only able to capture directional signalling due to the temporal ordering implemented by the EWC protocol. Despite these positive results, based on our observations of the standardised contrast, we find that TE-Full still provides the strongest discriminability between the absence and presence of signalling patterns. For this reason, we continue using TE-Full as a representative benchmark of directed FC in the subsequent analyses.

### Computational tractability of the EWC protocol

In this section, we compared the computational tractability of the EWC protocol by comparing full versions of TE and MI, and EWC versions of TE, cMI and the PC. We used these methods to estimate functional networks comprising increasing number of nodes. Nodal time series were sampled from source-localised MEG recordings (see Methods).

As expected, the computation time increased with network size in all cases. The conventional implementation of the TE was the most computationally intensive approach (Fig.3A), whereas PC-EWC estimation required ≈75% less time relative to TE-Full. However, the EWC implementation of TE was considerably quicker than the conventional approach and required just as long as the cMI-EWC. Interestingly, MI estimated from the full signal requires less time to compute than the closely related cMI-EWC. This is however due to an increase in the number of computations in the EWC approach, as we will discuss later.

In short, we find that both methods of estimation (Full or EWC) and the within-window EWC FC measure affect computation time. Importantly, using the EWC protocol generally leads to a decrease in computation time. In addition, the gain in computational time grows as a function of network size, indicating that EWC may confer pronounced benefits for the estimation fine-grained functional networks (e.g. >1000 nodes). Combined with our results from the previous section, we find that PC-EWC can be computed ≈4 times faster than TE-Full, while also resolving asymmetries in signalling.

### Inferring whole-brain interaction patterns from MEG recordings

Having compared both the accuracy and computational efficiency of the EWC to the conventional approach in-silico, in our final set of analyses, we extended the comparison onto empirical neuroimaging data. Specifically, we compute and compare PC-EWC and TE-Full from resting-state source-localised MEG recordings of 30 subjects, for the left hemisphere (right-hemisphere results Fig.S3). The temporal ordering was based on delays proportional to the inter-regional Euclidean distance (see Methods).

For each subject, we computed the correlation between the PC-EWC and TE-Full matrices and found that the two measures led to highly correlated estimates of functional connectivity (Fig 4B). The median correlation coefficient across subjects was r≈0.82 (p<0.0001) (Fig 4C top). Critically, although PC-EWC and TE-Full were very similar, the EWC protocol offered a speedup per epoch by a factor of ≈6.5 (Fig.4D bottom). In addition to the high subject-level agreement, the EWC implementation also allows us to study how signalling directions and strengths change across the scan, by estimating the degree of asymmetry and variance of estimates at different scales (event-, epoch-, and subject-levels) (Fig.S4).

**Figure 4.**
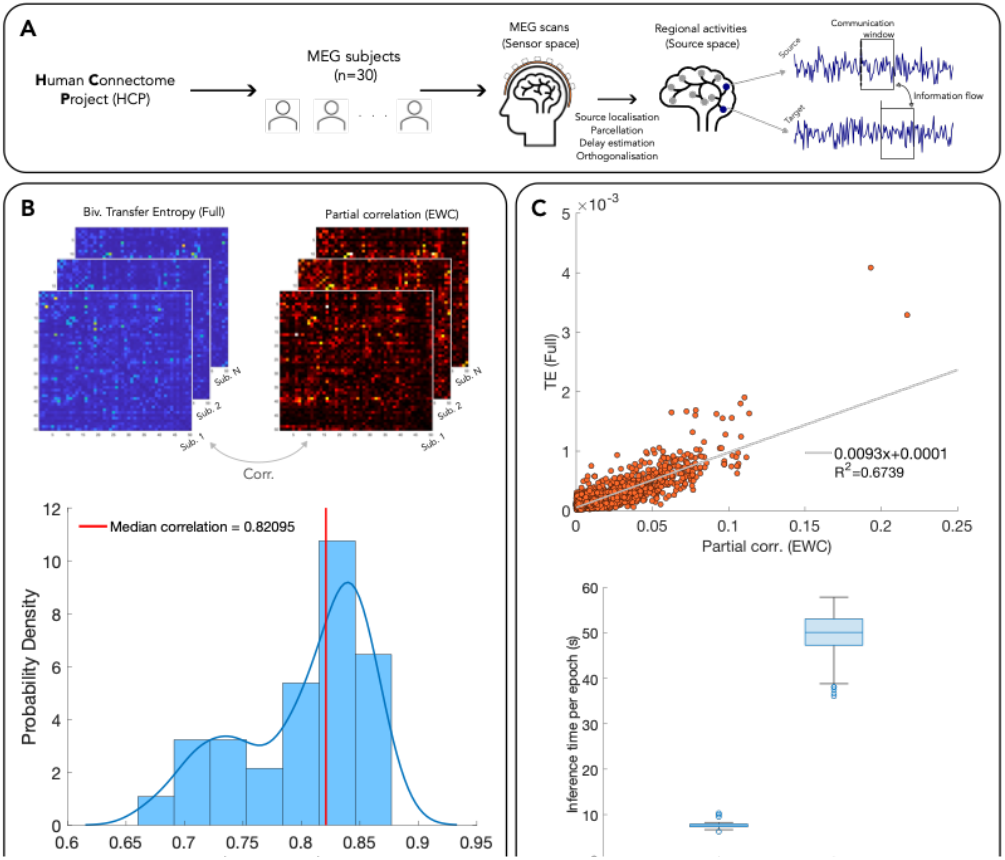
Application of EWC on source-localised MEG recordings. (A) Resting-state MEG scans of 30 subjects from the Human Connectome Project (HCP) were pre-processed, source localised, orthogonalized, and epoched into 10 second segments. (B) (Top) The subject-level FC for the left hemisphere was estimated as PC-EWC and TE-Full. (Bottom) Edgewise correlation distribution between the subject-level FC matrices. The red line marks the median correlation. (C) (Top) Scatterplot of edge weights for a representative subject (closest to the median correlation). Correlation aber excluding outliers (> 4 standard deviation) - r≈0.82, p<0.0001. (Bottom) inference time per epoch across subjects, for each of the methods.

To summarise, in this section we demonstrate a real-world application of the EWC, using PC-EWC and TE-Full. We show that the FC estimated as per the EWC shows good agreement with our conventional reference using the TE, and does so at a fraction of the computational cost. This lends support to the EWC as a viable technique for practical applications, particularly involving multiple functional components.

## Discussion

Understanding how the brain routes information is an open problem in neuroscience with crucial implications to our knowledge of perception and cognition. In this work, we focus on the first step towards addressing this problem – reliably inferring communication from neuroimaging data. We introduce a time series analytical technique, termed Event-marked Windowed Communication (EWC), to capture dynamic statistical relationships driven by inter-regional communication. This method quantifies inter-regional relationships using functional connectivity (FC) measures estimated on select subsamples of the signal. We demonstrated the merits and limitations of this method in silico, using different FC measures, and compared it to established methods of inferring directional relationships (Fig.2). We then applied it to source-localised MEG data, where high correlation was evident between subject-level FC patterns derived using EWC and conventional approaches (Fig.4). Importantly, we also showed that the EWC protocol is computationally efficient (≈6.5 times faster per 10s-epoch when using the PC) (Fig.3,4C-bottom), and additionally capable of capturing changes in asymmetric relationships over the scan (Fig.S4).

Studying how activity or information flows between neural elements is crucial to our understanding of the mechanisms of communication. In addition to EWC, multiple previous works have explored this important topic from diverse standpoints, such as neuronal avalanches^26,27,31^, direct stimulation-response measurements^33^, functional hierarchies^21^, FC dynamics^32,34^, and information-theoretic methods^25,35,36^, to name a few. EWC differs from previous work primarily in its handling of time series. As opposed to measures/features computed using the entirety of recordings or parts of them (as in sliding window approaches), EWC restricts FC estimation to select subsamples of the signal, to infer activity propagation between neural elements. By targeting the FC estimation to select subsamples of the scan, EWC effectively computes a resting-state “event-related potential”. Importantly, being a general and flexible framework, the EWC implementation can be easily incorporated into previously developed methodologies involving neural time series e.g. using feature vectors instead of raw time series^35^.

Estimating EWC comprises three steps: 1) event identification 2) temporal ordering 3) windowing/subsampling. Each of these elements have associated advantages and limitations. We discuss these points below.

Typically, FC is estimated for the entire scan, or in the case of dynamical-FC, for consecutive windows of the scan^24,37,38^. A limitation of such an approach to capture activity propagation is that inter-regional communication may occur only at select time epochs and this information may be obscured by internal dynamics. The purpose of the event-identification step in the EWC is to identify salient features in regional dynamics, that then serve as a reference point from which downstream effects are gauged. In task-based paradigms, these segments are generally marked by controlled stimulus triggers. On the other hand, in task-free recordings, change-point detection or point-process analysis-based tools can be used to capture salient changes in regional dynamics^39–42^. EWC reuses a simple method employed in previous works that traces activity propagation to parts of the signal that deviate from mean behaviour, through z-scoring^31^. Limiting the signal that is analysed to the proximity of the events additionally makes the EWC computationally tractable when compared to conventional approaches (Fig.3).

The temporal ordering aspect, which is designed to account for delays in signal conduction over the network, incorporates directionality into the protocol, irrespective of the FC measure used. This allows us to use undirected/symmetric measures like PC or cMI, to resolve transient asymmetric FC relationships (Fig.2B,C,E,F). Any symmetric FC measure (coherence, phase locking value, phase lag index etc.^43–45^) can similarly be rendered directional if implemented in this manner. Care must however be taken to ensure that the considered delays are within a reasonable range, to avoid capturing effects that might not likely be caused by the observed event.

Limiting the FC estimation to a window/subsample of the signal proximal to the events offers multiple advantages. It is well suited for scenarios where there is limited information regarding the spatiotemporal span of the effects, allowing us to capture extended effects as opposed to a local one (at a single time point). Furthermore, averaging the results over multiple windows (events) increases the robustness of the estimate. It also reduces the size of the signal for which the FC must be estimated, decreasing the computational time.

In contrast to conventional FC implementations however, EWC typically involves a higher number of number of computations – instead of a single computation per epoch, there are now computations associated with each event within the epoch (albeit involving shorter segments of data). This effect is apparent in Fig.3, where MI-Full is computed considerably faster than its EWC counterpart – cMI-EWC.

Although we find that there is a strong agreement between the subject-level PC-EWC and TE-Full, we noted that agreement in terms of their asymmetry or directionality was not statistically significant. This could be due to the PC and TE capturing different statistical dependencies in the data. This difference may have been subtle in the simple network-motif (resulting in both the PC and TE capturing the true signalling asymmetries), but more pronounced at larger scales.

### Methodological considerations

We designed EWC to capture communication-driven statistical relationships or as a statistical analogue of activity propagation. Its development involved the use of several simplifying assumptions to ensure computational and analytical tractability. These points need to be considered to fully appreciate the scope of this work and limitations. For example, a z-value-based event identification as we used in this study, will be sensitive to changes in the mean activity – this is evident in Fig.S2F, where an increase in the source firing rate results in firing events not being identified, resulting in diminished estimates. Similarly, errors in delay estimation, which affect the temporal ordering, can result in the effect of the event being missed entirely. The window length is also a crucial component and must be chosen so that it captures the immediate effects of the event, while also being long enough to ensure that there are enough datapoints for an accurate FC estimate. The window and epoch lengths also determine the resolution at which the dynamics of asymmetric relationships can be observed (event-/epoch-scale vs. subject-scale).

It is also important to remember that EWC is not a direct measure of communication but is instead a measure of the statistical consequence of communication. It can only be used to infer communication. Care must be taken while choosing the FC measure, since different measures capture different features of signal similarity (correlation, phase synchrony etc.), while also being valid only on certain derivatives of the original signal e.g. Mutual information can be computed between any signal (with different interpretations), whereas the phase lag index is computed between the instantaneous phase time series. The effect of FC measure choice is evident in Fig.2B,C,E,F, where the cMI indicates strong interaction from node 4 to 2, despite the lack of any interaction by construction. The PC on the other hand, does not capture this functional link. In the case of non-negative FC measures like the cMI, estimates can sometimes be inflated due a positive bias (despite filtering based on significance testing). This can be corrected by subtracting a null-derived estimate from the observed (significant) estimate.

Additionally, it must be noted that although our protocol restricts the estimation of the EWC to supra-threshold events, we do not claim that communication does not take place during the sub-threshold segments. The significant deviations are used to systematically reduce the analysis space, since it is more difficult to establish whether EWC estimates are due to communication as opposed to some other regional process or noise, in the sub-threshold segments.

## Conclusion

In conclusion, our work presents a new method to infer neural communication patterns through a restricted FC estimation approach based on activity propagation events. Our method yields FC estimates comparable to that of well-established techniques at a fraction of the computational time, opening up new avenues for the investigation of signalling in large networks of neural elements.

## Code availability

All the code used to generate activity propagation maps using the EWC is available at https://github.com/vmadanmohan/EWC.

## Methods

### Dataset

Resting state MEG scans of 30 subjects (22-35 years, 17F), along with associated MEG anatomical data, 3T structural MRI data, and empty-room recordings, were obtained from the Human Connectome Project (Van Essen et al., 2013), through the ConnectomeDB platform. The MEG scans varied in duration from 5-6 minutes, at a sampling rate (*SR*) of 2035Hz, and anti-aliasing low pass filtered at 400Hz.

### Processing

The MEG scans were processed entirely using the MATLAB^46^ -based Brainstorm software^47^, and in accordance with pipeline described in Brainstorm’s HCP-MEG tutorial^48^. MEG recordings were first coregistered to the subject’s structural MRI using the MEG anatomical data. A notch filter (60, 120, 180, 240 and 300Hz), followed by high pass filter (0.3Hz) were applied to resting state and empty-room recordings, to filter out power-supply and slow-wave/DC-offset artifacts respectively. Each subject’s recording was then visually inspected, along with the channel power spectral density, to weed out bad channels and bad time segments. The ECG and EOG recordings were then used to identify heartbeats and eye-blinks, after which associated artifacts were removed using their signal space projections (SSPs)^49^.

The source-level activities at 8000 points, defined by the fsLR4k mesh, were then estimated from the sensor-level recordings. This involved first computing the head model using overlapping spheres and constrained dipoles normal to the cortical surface, and estimating the noise covariance from the empty-room recordings. Source-level activities were then estimated using the dSPM method^50^ available in Brainstorm. The sources were then parcellated using the Schaefer-Yeo 7-network 100 atlas^51^, with the parcel activity computed as the mean of the constituent source activities.

The centroid coordinates of the Regions of Interest (ROIs) were used to define a Euclidean distance (ED) matrix between all possible pairs of regions. Assuming a conduction velocity of 10m/s, inter-regional distances were proportionally converted to time delays (*δ*) as

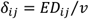

Where *v* is the conduction velocity (CV) of neural signals, set to 10 *m/s*.

Parcellated source-localised resting-state MEG recordings were corrected for source leakage effects by removing zero-lag correlations as per^52^, using the OHBA Software Library (OSL) package in Python^53^.

### Estimating inter-areal communication

#### “-EWC” implementation

The three main elements of EWC are – 1) event-identification, and 2) Temporal ordering based on conduction delays, and 3) estimation of the FC between event-marked windows/subsamples.

Source-level parcellated time series were first epoched into 10*s* segments. Epochs that contained bad segments were removed, and each epoch was then processed independently. The data within an epoch was first z-scored: This helped identify the spread of regional activities around their respective mean behaviours. For all regions, the timepoints at which the |*z*| >3 were labelled as “events” and marked strong significant deviations from mean behaviour. We argued that these significant events should cause measurable effects on the activity of the rest of the brain. Additionally, finite signal conduction times would imply that these downstream effects be delayed. We also focused the estimation of inter-areal communication to a segment of the time series starting at the event timepoint – termed a “communication window”, spanning a duration of 1*s*.

The following steps were executed iteratively, over all ROIs, for each event:

1. A region was chosen as a source, and a window of 1*s* was defined starting at the event. Any additional events that fell within this window were removed and not considered in the subsequent iterations, to avoid excessive computation. The window length was chosen to be short enough to capture the immediate effects of the communication events and minimise contributions from internal dynamics.
2. A window of similar length was placed at the time series of all other ROIs (targets) within the same hemisphere, at timepoints delayed in proportion to the Euclidean distance between the source and the target (see Processing). The 1*s* length of the window can also accommodate the effects of source signals travelling at velocities much slower than *10m/s* (used in the calculation of inter-regional delays).
3. The FC measure (PC, cMI) was computed between all pairs of the source and target signals within the respective windows, conditional upon the target’s immediate past activity over the duration of one window, up to the event. This conditioning was done to remove biases due to internal dynamics (not associated with communication). The FC estimates were checked for significance, with a Bonferroni-corrected threshold of 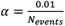, and set to zero for p-values above this threshold. The final measure is termed the Event-marked Windowed Communication (EWC).
4. The event-level EWC values from the source to all possible targets were then averaged to yield the source’s epoch-level EWC.

Once the above steps were completed for all possible sources, we were left with an epoch-level *N* × *N* EWC matrix, that captures the communication between all source-target pairs, averaged over all the events in the epoch. For a given subject whose scans were divided into *M*epochs, the communication protocol yields an *N* × *N* × *M* EWC matrix over the entire scan duration. This matrix is then averaged across epochs to give the subject-level EWC matrix.

#### “-Full” implementation

Source-level parcellated time series were first epoched into *10s* segments. Epochs that contained bad segments were removed, and each epoch was then processed independently. The FC was computed in a pairwise manner, between all regions, between the entire epoch time series. Each FC estimate was significance tested with a threshold of *α =0*.*0* 1. Like the EWC implementation, the pairwise FC was computed for all epochs, yielding an *N* × *N* × *M* matrix (for *M* epochs) over the entire scan duration.

### Partial correlation

The partial correlation (PC) quantifies the linear association between two random variables, while discounting the effects of a third variable. In the EWC implementation, we set the third variable as the target’s past over a single window duration, up to the timepoint of the event. We compute the PC using MATLAB’s “partialcorr” function. The significance level was set to 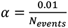. The partial correlation is a continuous function varying from -1 to 1, indicating perfectly negative to perfectly positive linear correlation respectively.

### Conditional Mutual Information

The conditional Mutual Information (cMI) is an information theoretic measure that quantifies the shared information between random variables (representing regional activities) in the context of the activity of some exogenous additional regions (represented as a conditioning variable). It is a variant of the widely used Mutual Information (MI)^25,54–56^.

Given random variables X and Y, containing the activities of two regions, and a third random variable Z, containing the activity of a third region (or a set of multiple regions), the cMI is measured in terms of entropies as:

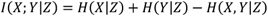

Where *H*(*X*|*Z*)and *H*(*Y*|*Z*)are the conditional entropies associated with variables X and Y, and *H*(*X,Y*|*Z*)is the conditional joint entropy.

The cMI is a symmetric measure, that varies from 0 when the two random variables are independent of each other, to *∞*when they are identical.

The target’s past activity was used as the conditional variable, to remove any biases to the measured communication from internal dynamics.

### Transfer Entropy

The Transfer Entropy is a widely used information-theoretic measure of directional influences ^2,3,25,36,57,58^. Specifically, for two random variables X and Y, it quantifies how much uncertainty in the future of Y (target) is reduced through the knowledge of the past of X (source), conditional on the past of Y. It is technically a special case of the cMI, where the shared information is estimated between relatively time-shifted random variables:

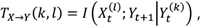

Where 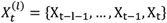. There are multivariate variants of the TE, which also condition it on other possibly mediating variables. In this work, we use its simplest form – the bivariate TE. Multivariate forms of the TE are generally better than other techniques at capturing ground truth asymmetries but are considerably more data- and computation-intensive^25^.

We used the Gaussian estimator in the Java Information Dynamics Toolkit (JIDT)^59^ to compute all information theoretic quantities. Using the Gaussian estimator allowed each estimate to be significance-tested analytically from the *χ*^2^-distribution. The estimators were run with default properties, except for the source-destination delay in the TE estimator, which was changed from the default value of 1, to the Euclidean distance-based inter-regional delay (see Processing). All information theoretic quantities (including the EWC) are measured in nats.

### Demonstration on a network model

In order to identify the merits and limitations of our approach prior to its application to neuroimaging data, we tested it on the activities of a four-node system with Linear Stochastic Model^60^ dynamics. The dynamics of a node *i* is described as:

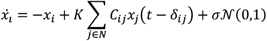

Where *C*_*ij*_is the connectivity strength between nodes *i* and *j, K* is a global coupling parameter (*K=* 1), *δ*_*ij*_ is the time delay between *i* and *j* (proportional to the Euclidean distance), 𝒩(0, 1) denotes random standard normal noise, and *σ* is the noise amplitude. Three of the nodes were connected in a linear chain-like topology, and the fourth node was isolated from the rest of the system i.e. *C*_12_*= C*_21_*= C*_2*3*_ *= C*_32_*=* 1; *C*_1*3*_ *= C*_31_*=* 0; *C*_4*i*_ *= C*_*i*4_ *=* 0 ∀*i* ∈ {1,2,3}.

In addition to the LSM dynamics, a Poisson process caused both the terminal nodes to “pulse” randomly, with an average frequency *v*, and with a pulse amplitude of 0. 1. This resulted in the dynamics of the central node to be a mix of its internal dynamics, and random inputs from the two terminal nodes. While Poisson spiking is a convenient means of exercising control over ground-truth signalling, it can result in an inflation of nodes’ mean activities from the “baseline” at high *v*, due to rapid fluctuations.

The communication protocol was tested across systematic variations of *σ, δ*_*23*,_ and *v*.

For each set of parameter values, we simulated the dynamics for *205s*, at a sampling rate of 2035*HZ*, using an Euler integrator. Epoch duration and communication window durations were set to *10s* and 1*s* respectively. 20 trials of each simulation were carried out.

Communication was inferred using the TE (Full implementation), cMI and PC (EWC implementation) (see Protocol), on the activity time series of all four nodes.

### Computation time benchmarking

To compare the computation times for different protocols as a function of network size, activity propagation was inferred as TE-Full, MI-Full, TE-EWC, cMI-EWC, and PC-EWC for time-series data of 200s. To ensure that the estimated computation times were realistic, the recordings were randomly sampled source localised MEG recordings (see Processing). The following steps were repeated for a network size *N* (for 10 trials):

1. *N* regions are randomly chosen, and the associated regional time series were trimmed to 200s (sampling frequency = 2035Hz).
2. Inter-regional delays were set to 0.015ms.
3. Activity propagation was estimated between all pairs of regions as TE-Full, MI-Full, TE-EWC, cMI-EWC, and PC-EWC.
4. Repeat with a new random selection of regions.

## Acknowledgments

V.M.M. is supported by the Melbourne Research Scholarship, University of Melbourne. R.F.H.C. is funded by a NHMRC Emerging Leadership Investigator Grant (Grant number: 2017527). C.S. acknowledges support from the Australian Research Council (Grant number: DP170101815). A.Z. is supported by an ARC Future Fellowship and the Rebecca L. Cooper Foundation. Data were provided [in part] by the Human Connectome Project, WU-Minn Consortium (Principal Investigators: David Van Essen and Kamil Ugurbil; 1U54MH091657) funded by the 16 NIH Institutes and Centers that support the NIH Blueprint for Neuroscience Research; and by the McDonnell Center for Systems Neuroscience at Washington University. This research was supported by The University of Melbourne’s Research Computing Services and the Petascale Campus Initiative.

