## Supplementary Information for "Event-marked Windowed Communication: Inferring activity propagation from neural time series"

## S1.

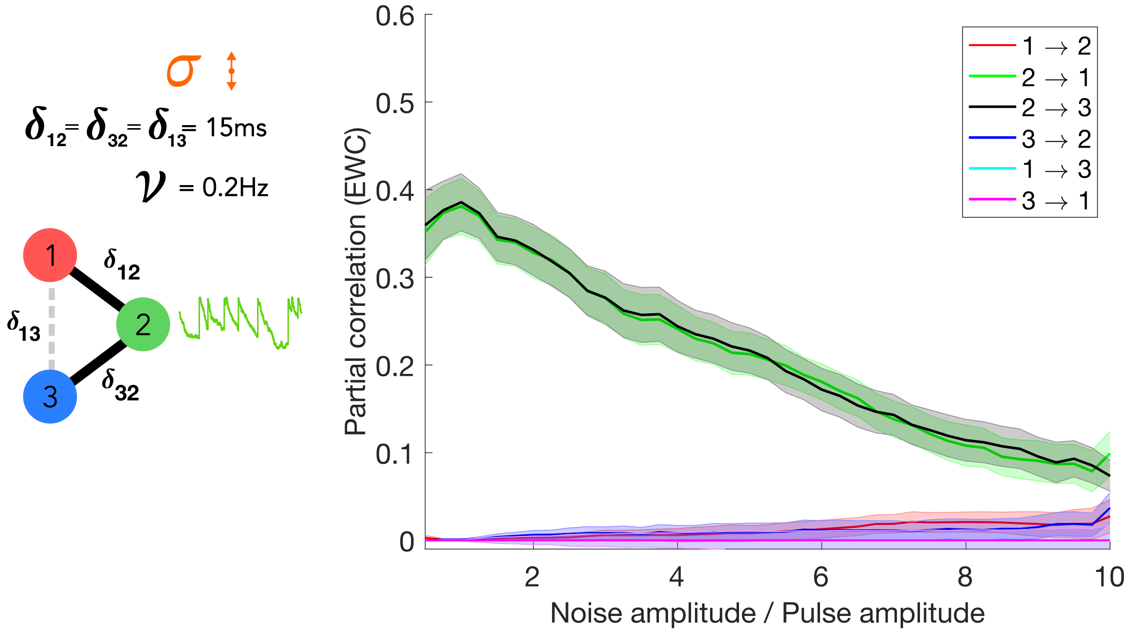


Figure S1 - Communication over a network motif with a common source (node 2). Nodes 1 and 3 follow LSM dynamics, whereas node 2 follows LSM dynamics in addition to Poisson process firing at a mean frequency of 0.2Hz. We observe that despite the equidistant targets (nodes 1 and 3) driven by a common source (node 2), the EWC method ascribes negligible communication from 1 → 3 and 3 → 1.

## S2.

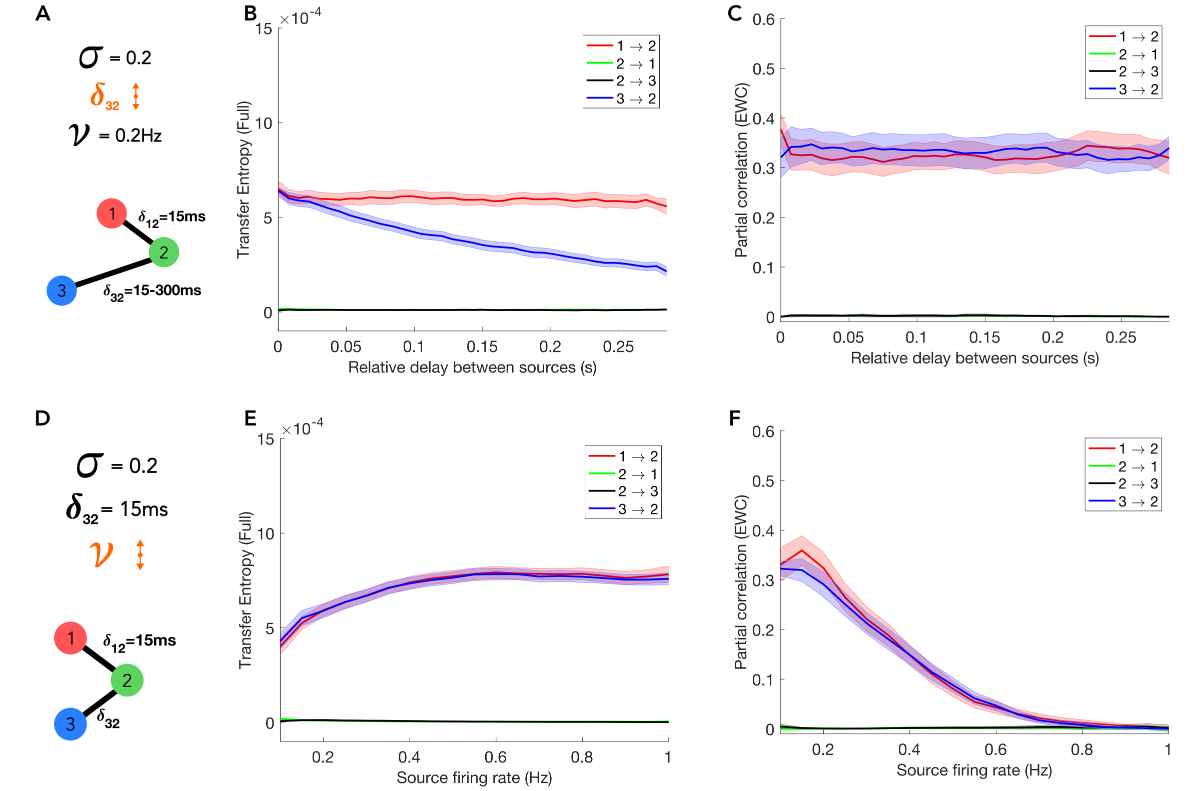


Figure S2 - Communication over a network motif with Linear stochastic model dynamics, with an addition of a Poisson spiking process in the dynamics of nodes 1 and 3. The delay between nodes 1 and 2 is fixed at 15ms for all simulations. We compared TE-Full and PC-EWC, over a range of parameters. (A-C) Conduction delay (distance) variation – The delay between nodes 3 and 2 is varied from 15ms to 300ms, whereas the noise amplitude and source firing rates are set at 0.2 and 0.2Hz respectively. We find that PC-EWC performs consistently over all delays, whereas the magnitude of TE-Full falls with increasing delay. (D-F) Source firing rate variation – The average firing rate of the sources (nodes 1 and 3) is varied from 0.2Hz to 1Hz, the noise amplitude is set to 0.2, and delay between nodes 3 and 2 is set to 15ms. We observe that the PC-EWC's performance drops quickly with an increase of firing rate. The TE shows an increase in communication strength with the source firing rate and plateaus around 0.6Hz. Plots are maximally smoothed (i.e. using all datapoints for a parameter value) to reveal trends. All simulations were repeated for 10 trials. Shading = ±SEM.

16

## S3.

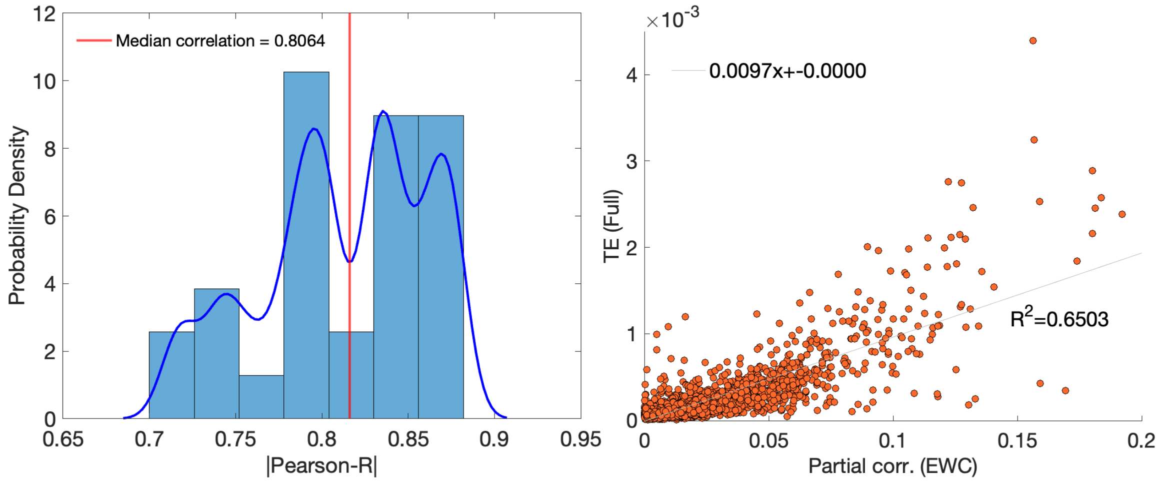


Figure S3 - The subject-level FC for the right hemisphere was estimated as PC-EWC and TE-Full. (Left) Edgewise correlation distribution between the subject-level FC matrices. The red line marks the median correlation. (Right) Scatterplot of edge weights for a representative subject (closest to the median correlation).

17

## S4.

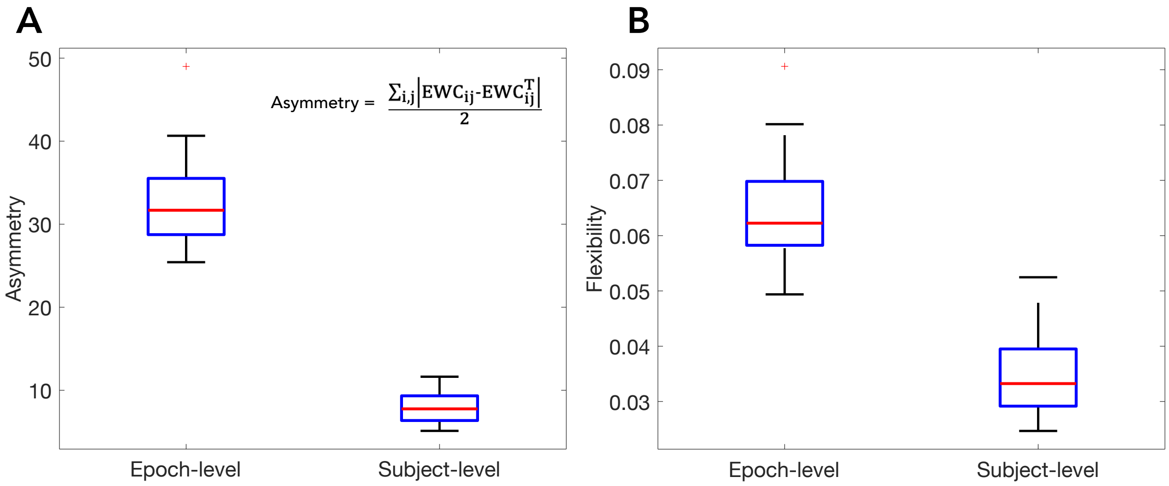


Figure S4 - Distribution of asymmetry and flexibility values of PC-EWC patterns at the epoch- (averaged over events) and subject-level (averaged over epochs). **(A)** (Inset) The asymmetry is defined as the halved difference between upper and lower triangle elements of the PC-EWC matrices. We observe that epoch-level PC-EWC matrices are highly asymmetric, indicative of unidirectional information routing, in contrast to the subject-level matrices, which show greater symmetry, indicative of bidirectional information routing. **(B)** The variability of PC-EWC values (quantified by the standard deviation (SD)) at different time-scales captures how dynamic or flexbile communication patterns are. We observe that average epoch-level flexibility (variability of EWC across events) is higher than the average flexibility at the subject-level (variability across epochs).

18
